# Estimating the abundance of benthic macrofauna from across shore transects

**DOI:** 10.1101/542258

**Authors:** Stephen L. Totterman

## Abstract

Benthic macrofauna of the intertidal zone are commonly sampled at low tide by excavating and sieving quadrats or cores on across shore transects. This paper examines different equations for estimating transect scale abundance, introduces the sampling theorem for calculating the maximum sampling interval and uses simulations to investigate sampling issues. Adaptive transect designs are recommended, where the sampling interval and replication of quadrats are carefully selected according to the width and density of macrofauna bands.

## INTRODUCTION

Sandy beach benthic macrofauna commonly aggregate in discontinuous ‘belts’ or ‘bands’with unimodal across-shore distributions (*e.g.* Defeo and Rueda, 2002). Across shore transects for sampling benthic macrofauna typically extend from the high water mark to the low water mark and one or more quadrats are sampled at each sampling ‘level’ within a transect (Figure 1). The terms quadrat (usually a square ‘box’) and core (a cylinder) are exchangeable in this paper: both refer to a sampling device with fixed area and depth. Fixed interval transects have a fixed level interval (*i.e.* along-slope spacing between adjacent levels) and the number of levels varies with beach width (transect length). Fixed area designs vary the level interval in proportion to transect length so that the number of levels and transect area are constant (Schoeman *et al.*, 2003).

**Figure 1.**
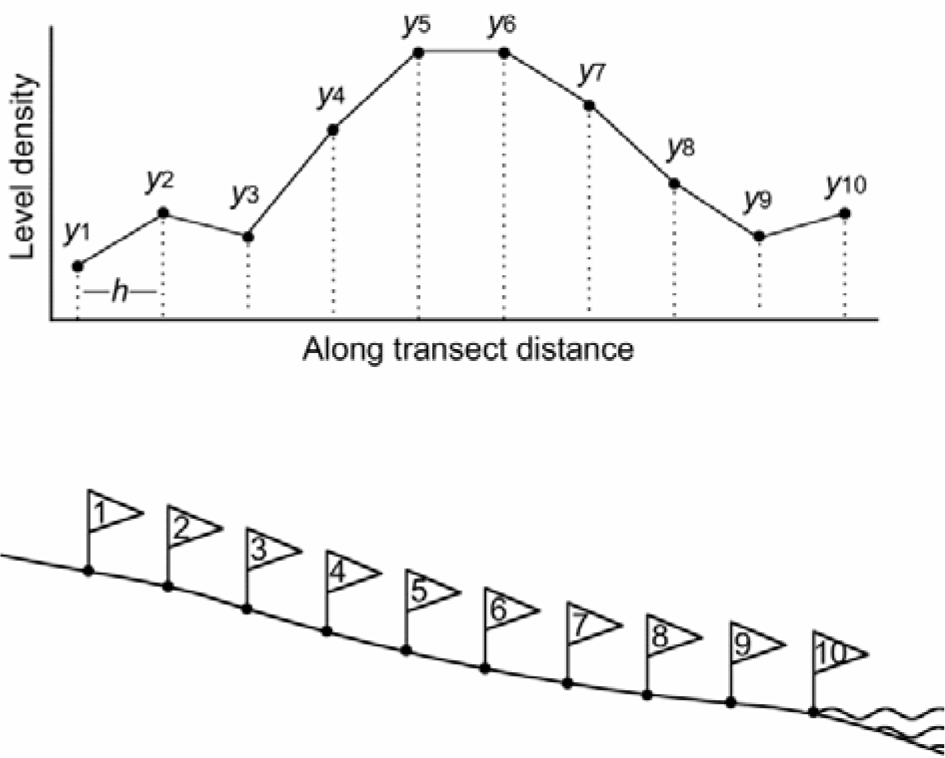
Illustration of an across shore transect with 10 intervals (lower) and the corresponding density-distance plot for benthic macrofauna recovered (upper) with linear interpolation trapezoid boundaries indicated by dotted lines. The fixed interval width is *h* and the level densities are *y*_*i*_.

Transect scale counts (*i.e.* the sum of quadrat counts over a transect) increase as more quadrats and larger quadrats are sampled. Counts can be standardised as areal densities (individuals/m^2^). Mean level density 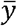 is (Figure 1):

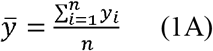

where *y*_*i*_ is the density (individuals/m^2^) for level *i* and *n* is the number of levels sampled. The level area (number of quadrats per level × individual quadrat area) is often fixed and then 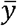 is simply the transect count divided by the transect area:

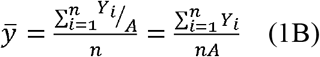

where *Y*_*i*_ is the count for level *i* and *A* is the constant level area. For fixed area designs, the transect area *nA* is constant and one could just as well report transect counts (individuals/transect).

Mean level density (Equation 1) doesn’t take into account beach width and the across shore distribution of levels. The standard solution for these problems is linear interpolation and integration of level densities along the transect length (Brown and McLachlan, 1980). The resulting linear density *LD* (individuals per along shore metre of beach) is the area under the density profile (*i.e.* the sum of trapezoids in Figure 1):

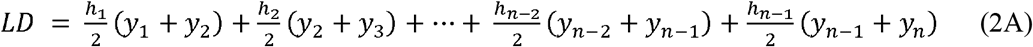

where *h*_*i*_ is the interval between levels *i* and *i* + 1. For fixed interval transects:

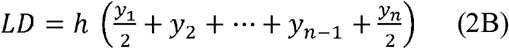

where *h* is the fixed level interval.

The simple individuals per strip transect (*IST*; individuals/m) equation (Defeo 1996) has been popular for calculating linear density. *IST* is the product of transect length and mean level density:

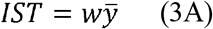

where *w* is the transect length. For fixed interval transects:

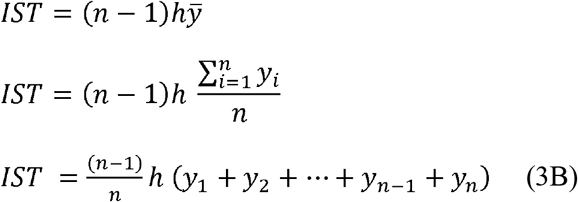

where *h* is the constant level interval and (*n* – 1)*h* is the transect length (Figure 1). This fixed interval *IST* (Equation 3B) is not identical to integrated density (Equation 2B). A positive bias in *IST* can result from the larger weighting of the terminal densities *y*_1_ and *y*_n_. A negative bias can result from the coefficient (*n* – 1)/*n*. *IST* and integrated density are strictly equivalent only when *h* is constant, *y*_1_ = *y*_n_ = 0 and for *n* → ∞. Relaxing these conditions, *IST* and integrated density are approximately equal when *h* is constant, 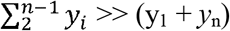 and when *n* is large. It is dangerous to assume that the positive and negative biases approximately cancel out.

Schoeman *et al.* (2003) studied the effect of transect level interval width on total *IST* estimates and accidentally provided some examples of bias using the *IST* equation 3A. They sampled six transects on three beaches in the Eastern Cape region of South Africa. The bivalve *Donax serra* was dominant in transects from Sundays and Maitlands beaches, with 34–61% of total counts. *D. serra* aggregations are typically found in the mid to upper intertidal zone on Eastern Cape beaches (Donn *et al.* 1986) and it can be assumed that total counts for Sundays and Maitlands beaches decreased strongly towards the top and bottom transect levels. Positive bias in *IST* would then have been small and the results of Schoeman *et al.* (2003) do show negative bias close to that expected from Equation 3B (Figure 2).

**Figure 2.**
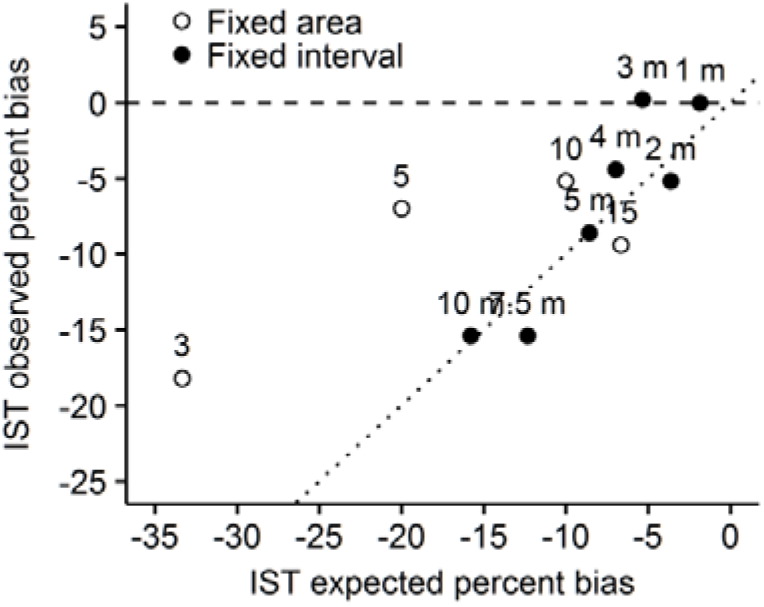
Mean observed bias and expected bias for fixed interval (black fill: 1, 2, 3, 4, 5, 7.5 and 15 m) and fixed area (white fill: 3, 5, 10 and 15 levels) individuals per strip transect (*IST*) estimates for Sundays and Maitlands beaches from Schoeman *et al.* (2003). *IST* is negatively biased when the level interval is wide and the number of levels is small. The dotted line *y* = *x* indicates perfect agreement. Observed bias results are means from Figure 2 in Schoeman *et al.* (2003). Expected bias assumes the top and bottom level densities are zero (Equation 3B). For fixed interval results, transect lengths (number of quadrats × quadrat edge length) were calculated from Table 2 in Schoeman *et al.* (2003) and then used to calculate the number of transect levels and expected bias. Undersampling is suggested for three and five level transects in the Discussion.

Given realistic parameters, simulation studies can help understand and improve sampling designs at lower cost than field studies (Miller and Ambrose, 2000). The remainder of this paper uses simulations to investigate transect sampling issues.

## METHODS

Single macrofauna bands were simulated with a Gaussian density function. This unimodal ‘bell-shaped’ curve is similar to the across shore distributions of sandy beach macrofauna measured in the field (*e.g.* Defeo and Rueda, 2002). Simulations varied the beach with (transect length = 30, 60, 120) and true abundance (total count = 100, 200, 400), location (mean = 0.5, 0.75 and 0.9 × length) and scale (*sd* = 1, 2, 5, 10, 20) of the density function. The widths of simulated bands, calculated from the lower 2.5% to upper 97.5% quantiles of the Gaussian curve, were 4, 8, 20, 40 and 80.

Quadrat scale patchiness in benthic macrofauna is a well-known sampling issue (Andrew and Mapstone, 1987; James and Fairweather, 1996; Defeo and Rueda, 2002; Defeo and McLachlan, 2005) and this ‘noise’ tends to obscure unimodal distributions in field data. Among-quadrat variability in counts within transect levels was simulated with a negative binomial count model. The expected value (mean) is set by the Gaussian density curve above and the variance is (Venables and Ripley, 2002):

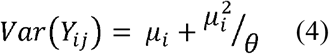

where *Y*_*i*_ is the count for quadrat *j* at level *i* and *μ*_*i*_ is the true mean. The shape parameter □ describes overdispersion relative to the poisson count model. Different macrofauna species can show spatial patterns varying from ‘random’ (□ ∼ ∞) to ‘clumped’ (□ → 0) and □ is scale-dependent, being affected by the quadrat size and spacing between replicate quadrats (Andrew and Mapstone, 1987). Pilot studies are needed to □ for the macrofauna species of interest as well as to estimate the width of bands. Here, □ = 1.6 is based on extensive sampling of the beach clam *Donax deltoides* using 0.1 m^2^ quadrats with 5 paces between quadrats (56 clam bands with > 0.5 clams/m^2^ mean density on 15 beaches, each sampled with 7 quadrats; unpublished data).

There were three simulations: 1) fixed interval sampling of level densities (*i.e.* noise free data) at intervals 1, 3, 6, 10, 20 and 40; 2) fixed area sampling of level densities with 3, 6, 11 and 16 equally-spaced levels; 3) fixed interval sampling of quadrat counts (*i.e.* noisy data) with 1, 3 and 10 replicate 0.1 m^2^ quadrats per transect level.

Simulation code for *R* version 3.4.4 (R Core Team, 2018) is provided in the Supplementary Material.

## RESULTS

Density sampling simulations demonstrated two sources of bias for integrated transect density (Equation 2): low sampling frequency and truncation (examples in Figure 3). For transects without truncation, Figure 4A shows that the transect interval and, therefore, the number of transect levels (or, more broadly, ‘sampling effort’) is a poor predictor of bias. Under sampled transects can either under-(Figure 3A) or over-estimate (Figure 3B) abundance. Figure 4B demonstrates the well-known sampling theorem (Meijering, 2002): bias is negligible when the sampling interval is less than 0.5× the width of the band. This critical sampling interval is commonly known as the Nyquist interval. Figures 3C and 4C show that, for transects with adequate sampling frequency, integrated densities are negatively biased according to the proportion of the band truncated.

**Figure 3.**
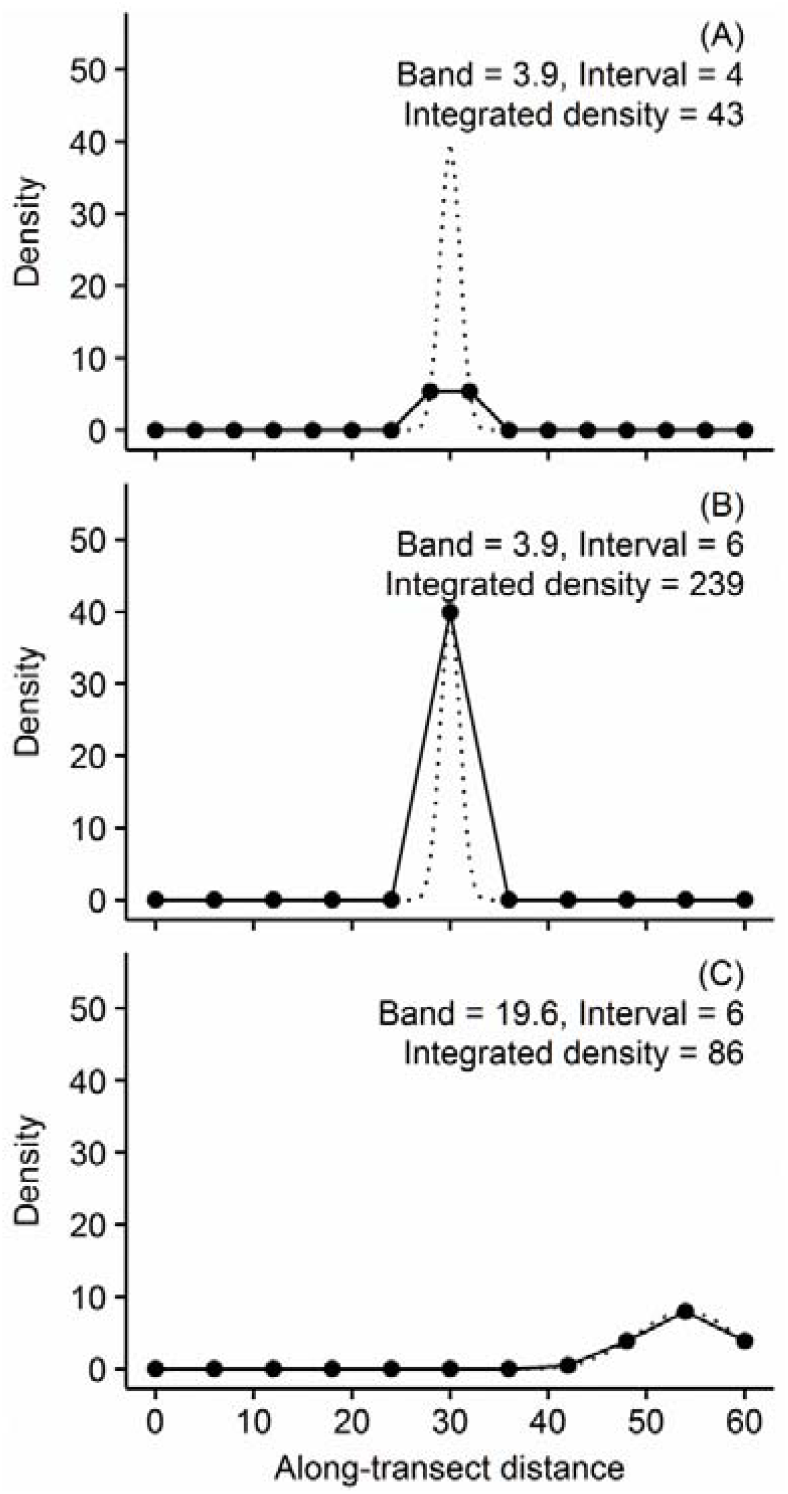
Fixed interval density sampling simulation examples of negative (A) and positive (B) undersampling bias and truncation bias (C). The true abundance was 100. Simulated densities are shown as dotted lines. Sampled densities are plotted as solid circles and solid lines show linear interpolation.

**Figure 4.**
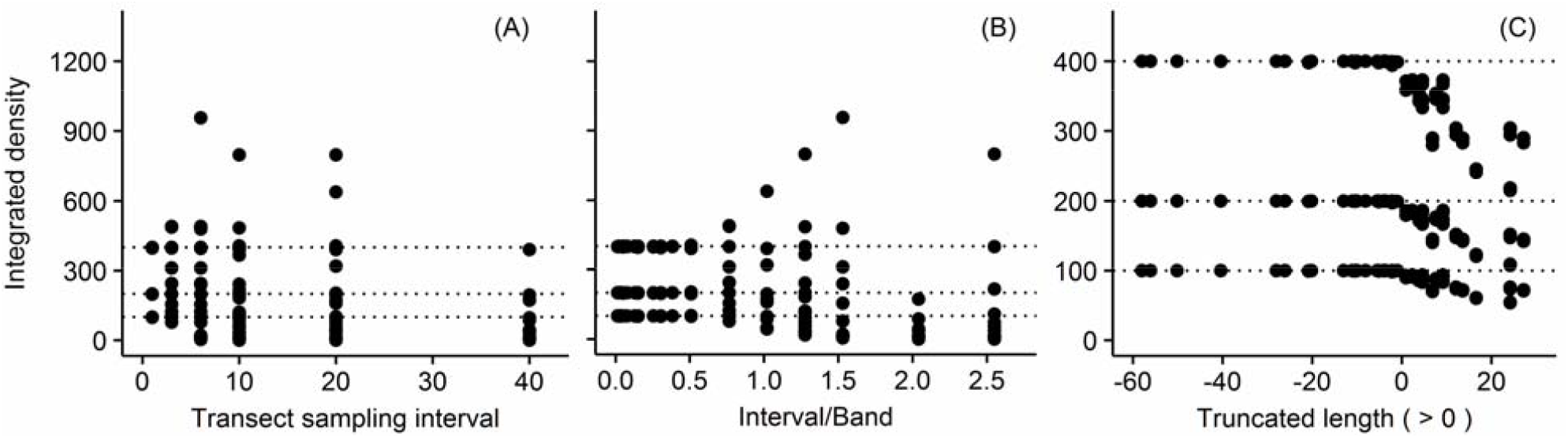
Fixed interval density sampling simulation results for various transect sampling intervals (A and B; without truncated distributions) and various truncation extents (C; without undersampling). True abundances are shown as dotted lines (100, 200, 400). Integrated densities were unbiased when the sampling interval was less than 0.5× the band width (B) and when the band was not truncated (C).

For density sampling simulations with adequate sampling frequency and no truncation, Figure 5 compares integrated transect density (Equation 2B), *IST* (Equation 3B) and mean level density (Equation 1A) with true abundance. *IST* was negatively biased when the number of levels is small (Figure 5B). Mean level density was approximately proportional to true abundance only when beach width (transect length) was constant (Figure 5C). Figure 6 shows that mean level density is approximately inversely proportional to transect length. For example, when the beach is wide and the band is relatively narrow, there are many zero level densities outside of the band and the mean is low. The width of the band has little influence on mean level density because, for some constant total count, wider bands have lower level densities but also relatively few zero level densities compared to narrow bands.

**Figure 5.**
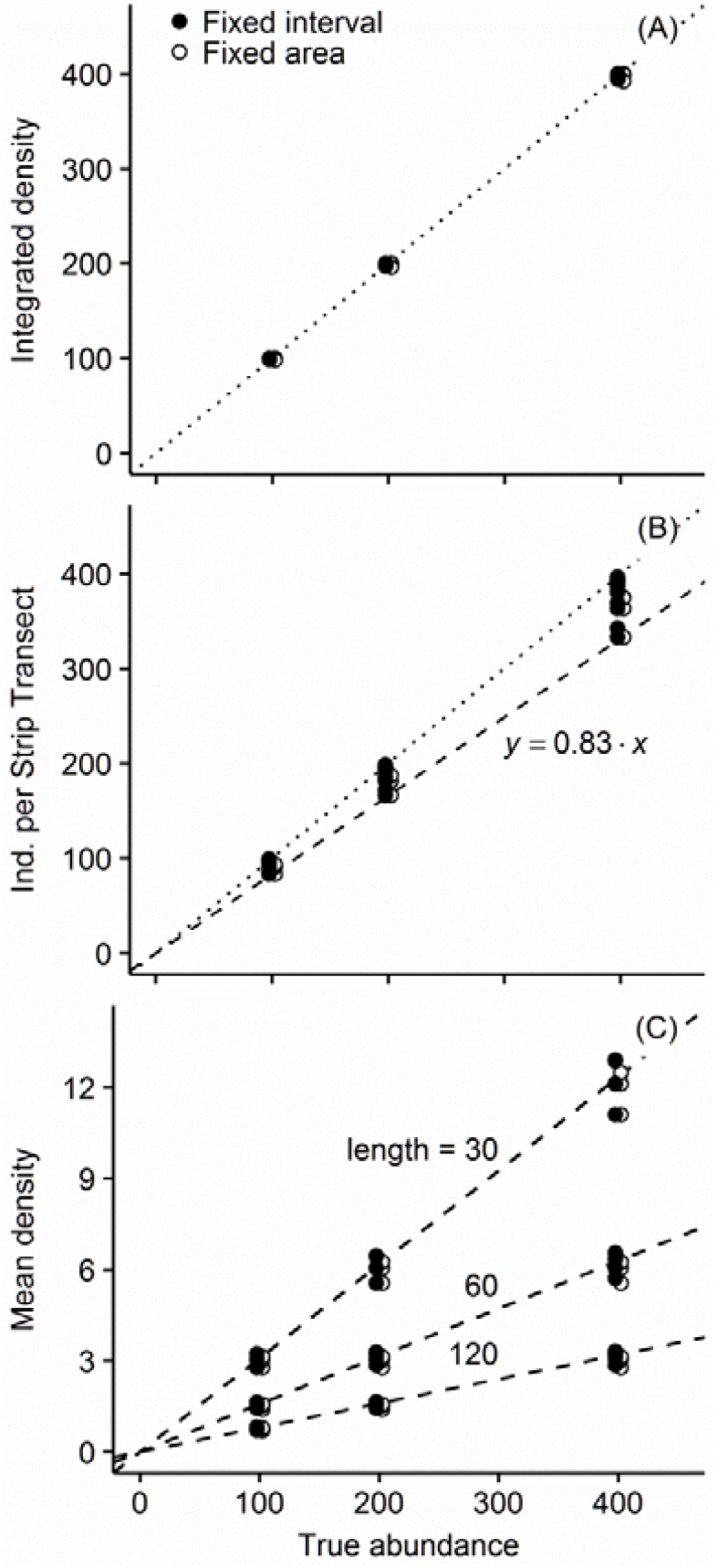
Comparisons of integrated density (A; Equation 2B), individuals per strip transect (B, Equation 3B) and mean density (C; Equation 1A) with true abundance for fixed interval (black fill) and fixed area (white fill) density sampling simulations. Results with truncation and undersampling have been removed. The dotted lines in A and B indicate perfect agreement. The dashed line in B indicates bias for six transect levels (slope = 5/6). The dashed lines in C correspond to transect lengths 30, 60 and 120. Fixed interval and fixed area points have been ‘jittered’ slightly along the *x*-axis to reduce overplotting.

**Figure 6.**
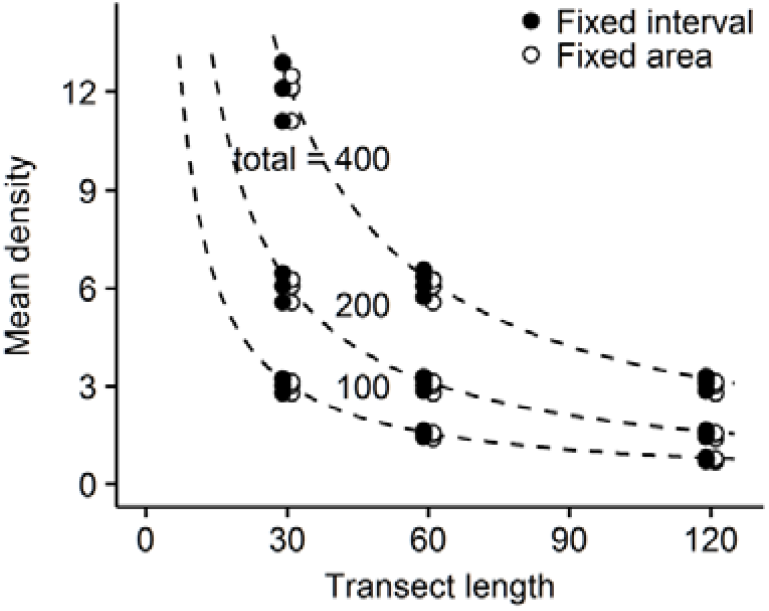
Mean density (Equation 1) versus transect length (beach width) for fixed interval (black fill) and fixed area (white fill) density sampling simulations. Results with truncation and undersampling have been removed. Dashed lines correspond to true abundances 400, 200 and 100. Fixed interval and fixed area points have been ‘jittered’ slightly along the *x*-axis to reduce overplotting.

For quadrat count sampling simulations with adequate sampling frequency and no truncation, Figure 7 compares integrated transect density (Equation 2B), *IST* (Equation 3B) and mean level density (Equation 1A) with true abundances. Noise tends to masks bias in *IST* and non-proportionality for mean level density. Noise increased for higher densities because of the assumed quadratic mean-variance relationship (Equation 4). Noise decreased with larger numbers of replicate quadrats per level because of the well-known relationship between the variance of a mean and the sample variance (Zar 1999):

**Figure 7.**
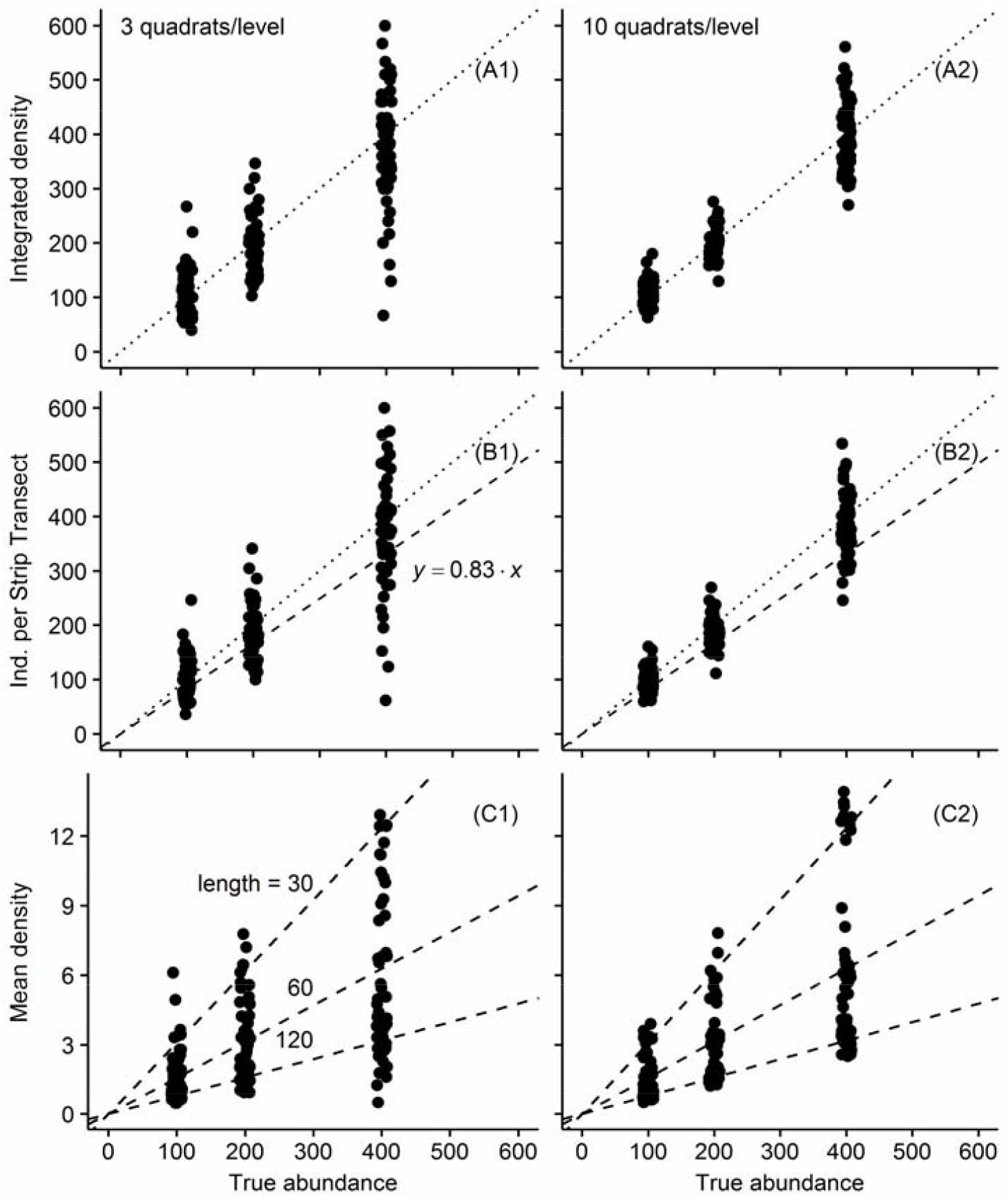
Comparisons of integrated density (A; Equation 2B), individuals per strip transect (B, Equation 3B) and mean density (C, Equation 1) with true abundance for count simulations of single fixed-interval transects with three (1) or ten (2) replicate quadrats per transect level. Results with truncation and undersampling have been removed. The dotted lines *y* = *x* in A and B indicate perfect agreement. The dashed line in B indicates bias for six transect levels (slope = 5/6). The dashed lines in C correspond to transect lengths 30, 60 and 120. A small amount of random noise (‘jitter’) has been added to the totals (*x*-axis) to more clearly show the distribution of densities. These results can be compared with noise-free results in Figure 5.

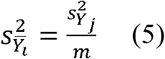

where 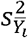 is the variance of the level mean count 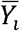 and 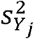 is the variance among the *m* replicate quadrat counts *Yj*. Over sampling transect levels similarly reduces variance because level densities sum quadrat counts and then transect densities sum level densities, similar to increasing the number of replicate quadrats (Figure 8).

**Figure 8.**
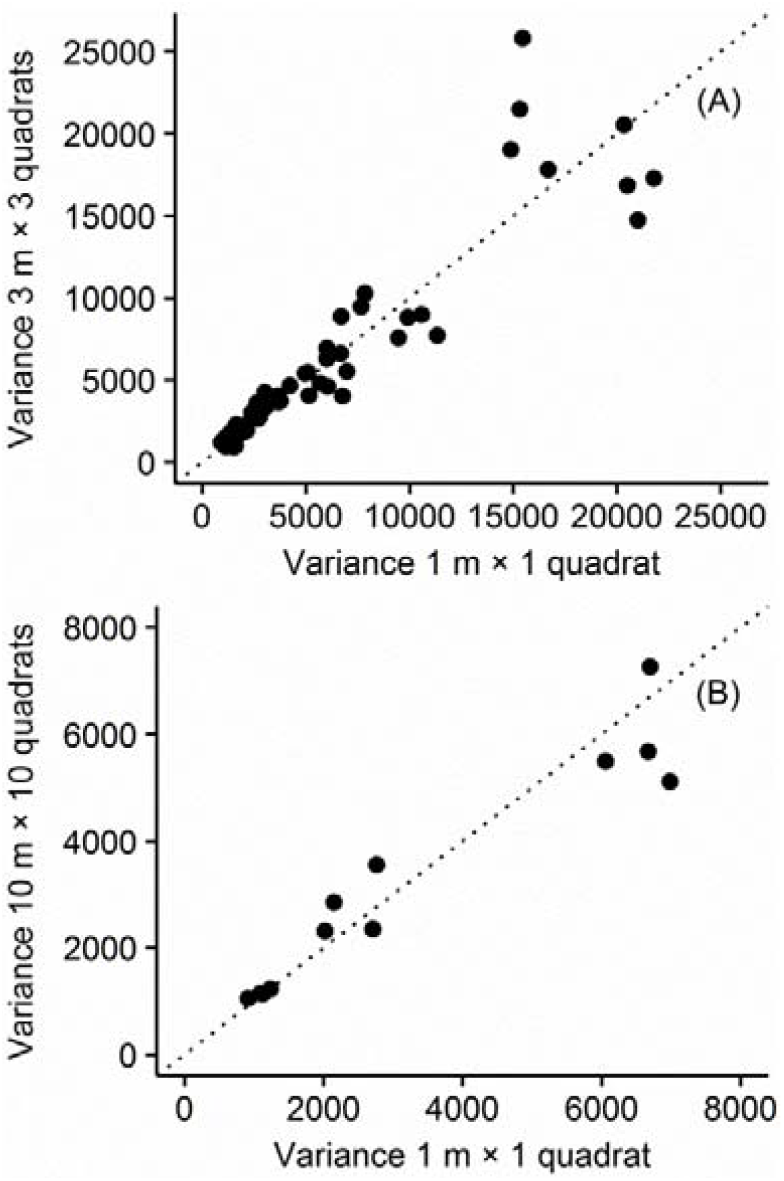
Comparisons of integrated density variance for quadrats averaged across shore (oversampling) and quadrats averaged alongshore (replication of quadrats within levels) for count simulations. Variances are computed from fixed-interval transects after removing truncated and undersampled results. The difference in sampling effort (number of levels × quadrats/level) was consistently +2 quadrats (greater effort with 3 replicates at 3 m intervals) in A and +9 quadrats (greater effort with 10 replicates at 10 m intervals) in B. The dotted lines *y* = *x* indicate perfect agreement.

## DISCUSSION

Two other recent efforts to improve across shore transect designs are Schoeman *et al.* (2003) and Schlacher *et al.* (2008). Schoeman *et al.* (2003) recommended a 3 m transect level interval, although that study used the wrong *IST* equation (Equation 3B). Nevertheless, erratic mean bias in their results for three and five level fixed area results suggests undersampling and agreement between observed and expected bias in their results for *n* > 5 transect levels suggests adequate sampling frequency for intervals ≤ 10 m (Figure 2). The 3 m interval recommended by Schoeman *et al.* (2003) can result in oversampling.

A consensus of expert opinion in Schlacher *et al.* (2008), recommended 7–27 transect levels (increasing with beach width), with corresponding transect intervals of 3–10 m. Simulation results from this study showed that sampling with only three levels was not sufficient and that six levels was adequate only when the macrofauna band was wide relative to the transect length. The minimum seven levels recommended by Schlacher *et al.* (2008) seems reasonable but can result in undersampling of narrow bands.

The sampling theorem can help to avoid costly oversampling (sampling interval < critical) or dangerous undersampling (sampling interval > critical). To generalise the sampling theorem to multispecies sampling, the critical sampling interval is half the width of the narrowest species band. In practise, a larger interval can be used if the more abundant species occur in wider bands (*i.e.* species with lower abundance contribute little to the integrated total density and potential biases would also have a small effect). Effort can be efficiently allocated with adaptive level intervals: 1) a wide interval can be used to approximately delineate the band, and then 2) extra levels are sampled across the band to increase resolution where it counts. The integrated density equation (Equation 2A) can accommodate variable interval widths.

Next, researchers need to beware of truncation bias. Truncated sampled distributions are very common on sandy ocean beaches where quadrat sampling is difficult in the low tide swash zone and impossible in the near subtidal (Defeo and Rueda 2002). Although rarely acknowledged, studies that do not sample the entire across shore distribution must assume that the proportion sampled (and resulting negative bias) is constant for all transects. Consideration should then be given to some estimate of truncation bias and the magnitude of this bias relative to the effect sizes of interest.

Fixed area transects do not give mean level densities proportional to true abundance. The inverse relationship between mean level density and beach length could have been the motivation behind the *IST* equation 3. Nevertheless, the integrated density equation (Equation 2A) can be applied to fixed area transects with variable interval widths. One application for fixed area transects is when the researcher wants to consistently position transect levels relative to physical reference points, *e.g.* a seven-level transect from the drift line to the low tide swash line will consistently place level four at the mid-intertidal.

Noise tends to masks bias in *IST* and non-proportionality for mean level density. Thus few beach studies have recognised bias in mean level density (Equation 1) and *IST* (Equation 3) except Schoeman *et al.* (2003) where bias was estimated from the average of 100 resampled transects. Nevertheless, *IST* bias is easily avoided by using the more rigorous integrated density equation (Equation 2). Attention then turns towards variance in quadrat counts and level densities as a source of inaccuracy. Schlacher *et al.* (2008) suggested three replicate 0.1 m^2^ quadrats per level. Simulation results from this study showed substantial improvements in precision when increasing replication to 10 quadrats. Sampling effort can be efficiently allocated with adaptive numbers of quadrats per level: 1) a minimum of three quadrats are sampled at each level, and then 2) replication is increased across the macrofauna band where counts and count variances are higher. Oversampling levels can also reduce variance and results in Schoeman *et al.* (2003) did show that *IST* variance does decrease with narrower level intervals. However, it is simpler to keep the issues of undersampling bias (transect level interval) and count variance (replication of quadrats per level) separate when sampling transects.

## CONCLUSIONS

Modern across-shore transect designs often and unnecessarily use narrow, fixed level intervals (Schoeman *et al.* 2003). Narrow intervals can prevent *IST* bias (Equation 2), undersampling bias and increase precision. However, intensively sampled transects are costly and this can be a convenient excuse for sandy beach studies sampling only one or a few replicate transects (James and Fairweather, 1996).

Based on sampling theory and simulations, this paper advocates flexible and efficient transect designs where level intervals as well as numbers of quadrats per level are carefully varied according to the width and density of macrofauna bands. Graphs of the across shore distribution of level densities or counts should be presented so that readers can visually assess the adequacy of the transect design.

## Supporting information

Supplementary R simulation code

## ACKNOWLEDGEMENTS

This paper is unlikely to be published for two reasons: 1) conservatism in the field of Sandy Beach Ecology, and; 2) influence of authorities in this field over journal editors (*e.g.* O. Defeo and D. S. Schoeman would not recommend this paper).

## SUPPLEMENTARY MATERIAL

R code for simulating sampling of line transects and analysis (text file).

