## Supplementary R simulation code for "Estimating the abundance of benthic macrofauna from across shore transects"

# ---------------------------
# simulation 1 FIXED INTERVAL
# ---------------------------

# data
# ----
# 60 m is a typical transect length
# 100 is a typical abundance (for surf clams)
# sca has been carefully adjusted to be close to int*2
# 3 m is a recommended interval (schoeman et al. 2003) 

results<-matrix(nrow=3*3*3*5*6, ncol=14)
i<-0 # index
## configure across transect abundance and distribution
for (tot in c(100, 200, 400)){ # transect total (ind/m)
  for (len in c(30, 60, 120)) { # transect length
    for (loc in c(0.5*len, 0.75*len, 0.9*len)){ # band location (mean for dnorm)
      for (sca in c(1, 2, 5, 10, 20)){ # band scale (sd for dnorm)
        for (int in c(1, 3, 6, 10, 20, 40)){ # transect level interval
          i<-i+1
          results[i,1]<-tot
          results[i,2]<-len
          results[i,3]<-loc
          results[i,4]<-sca
          results[i,5]<-int
          ## configure sampling
          myx<-seq(0, len, int) # level locations
          ## check that len is divisible by int
          # forbidden combinations are (len, int) = (30, 20), (30, 40), (60, 40)
          if (max(myx)==len){
            ## sample
            myn<-length(myx) # number of levels, which also is the transect "cost"
            myy<-tot*dnorm(myx, loc, sca) # level densities (ind/m2)
            dens<-int*(0.5*myy[1]+sum(myy[2:(myn-1)])+0.5*myy[myn]) # integrated density
            band<-qnorm(0.975, loc, sca)-qnorm(0.025, loc, sca) # 95% band width
            trunc<-(loc+0.5*band)-len # positive truncation if upper limit of 95% band > length
            results[i,6]<-myn
            results[i,7]<-round(dens, 1) # integrated density
            results[i,8]<-results[i,7]-tot # bias
            results[i,9]<-round((dens-tot)/tot, 3) # relative bias
            results[i,10]<-round(mean(myy), 3) # mean level density
            results[i,11]<-round(band, 1)
            results[i,12]<-round(trunc, 1)
            results[i,13]<-round(trunc/band, 3) # relative truncation
            results[i,14]<-round(int/band, 3) # sampling frequency
          } # endif
        } # int
      } # sca
    } # loc
  } # len
} # tot
results<-as.data.frame(results)
names(results)<-c("tot", "len", "loc", "sca", "int", "levels", "ldens", "bias", "relb", "adens", "band", "trunc", "relt", "freq")

# result 1: sampling frequency problem
# ------------------------------------
# the critical sampling interval is 0.5 times the width of the band

# remove truncation
freqs<-subset(results, trunc<0)
# examine the effect of level interval and macrofauna band width on linear density
par(mfrow=c(2,2))
with(freqs, plot(ldens~int, ylim=c(0,1500)))
abline(h=100, lty="dotted")
abline(h=200, lty="dotted")
abline(h=400, lty="dotted")
with(freqs, plot(relb~int, ylim=c(-1,4)))
abline(h=0, lty="dotted")
with(freqs, plot(ldens~freq, xlab="int/band", xlim=c(0,3), ylim=c(0,1500)))
abline(h=100, lty="dotted")
abline(h=200, lty="dotted")
abline(h=400, lty="dotted")
abline(v=0.5, lty="dashed")
with(freqs, plot(relb~freq, xlab="int/band", xlim=c(0,3), ylim=c(-1,4)))
abline(h=0, lty="dotted")
abline(v=0.5, lty="dashed")
par(mfrow=c(1,1))

# result 2: truncation problem
# ----------------------------
# relative bias is proportional to relative truncation; e.g. if half the band is truncated, the integrated density is half

# remove undersampling
truncs<-subset(results, freq<=0.5)
# examine the effect of truncated distributions on linear density
par(mfrow=c(2,2))
with(truncs, plot(ldens~trunc))
abline(h=100, lty="dotted")
abline(h=200, lty="dotted")
abline(h=400, lty="dotted")
abline(v=0, lty="dashed")
with(truncs, plot(relb~trunc))
abline(h=0, lty="dotted")
with(truncs, plot(ldens~relt, xlab="trunc/band", xlim=c(-5,2)))
abline(h=100, lty="dotted")
abline(h=200, lty="dotted")
abline(h=400, lty="dotted")
abline(v=0, lty="dashed")
with(truncs, plot(relb~relt, xlab="trunc/band", xlim=c(-1,2)))
abline(h=0, lty="dotted")
abline(0,-1, lty="dashed")
text(0.7, -0.2, "y = -x")
par(mfrow=c(1,1))

# result 3: width problem
# -----------------------
# for fixed interval designs, mean level density is largely determined by transect length:
# 1) when the beach is wide, there are many low/zero level densities (i.e. outside of the band) and the mean is low
# 2) the width of the band has a relatively small influence because wider bands have low level densities but few low/zero level densities
# the first plot shows that integrated density is very close to the true density
# the fourth plot shows that IST is negatively biased for wider transect intervals (truncation removed and terminal level densities were zero) 

# remove truncation and undersampling
best<-subset(results, trunc<0) # no truncation
best<-subset(best, freq<=0.5) # no undersampling
# fit a model to adens ~ len
amodel<-lm(adens~I(1/len)*as.factor(tot), best)
summary(amodel)
# examine linear density, areal density and IST compared to totals and in relation to transect length
par(mfrow=c(2,2))
with(best, plot(ldens~as.numeric(tot)))
abline(0, 1, lty="dashed")
with(best, plot(adens~as.numeric(tot), col=len))
with(best, plot(adens~len, col=tot, xlim=c(10,120)))
xv<-seq(1, 120, 1)
yv<-coef(amodel)[1]+coef(amodel)[2]*(1/xv) # fitted tot = 100
lines(xv, yv)
yv<-sum(coef(amodel)[c(1,3)])+sum(coef(amodel)[c(2,5)])*(1/xv) # fitted tot = 100
lines(xv, yv)
yv<-sum(coef(amodel)[c(1,4)])+sum(coef(amodel)[c(2,6)])*(1/xv) # fitted tot = 100
lines(xv, yv)
with(best, plot(adens*len~as.numeric(tot), ylab="IST", col=int))
abline(0, 1, lty="dashed")
par(mfrow=c(1,1))

# evalute IST
with(best, plot(adens*len~tot, ylab="IST", col=levels))
abline(0, 1, lty="dotted")
# minimum acceptable levels was 6
abline(0, (6-1)/6, lty="dashed")
text(340, 250, "y = 0.83x")

# fit a model to adens ~ tot
# much the same as for fixed area results below
amodel2<-lm(adens~tot*as.factor(len), best)
summary(amodel2)

# -----------------------
# simulation 2 FIXED AREA
# -----------------------

# data
# ----
# 60 m is a typical transect length
# 100 is a typical abundance (for surf clams)
# 2, 6, 11, 16 levels is based on Schoeman et al. (2003) and adjusted to get integer valued intervals 

results2<-matrix(nrow=3*3*3*5*6, ncol=14)
i<-0 # index
## configure across transect abundance and distribution
for (tot in c(100, 200, 400)){ # transect total (ind/m)
  for (len in c(30, 60, 120)) { # transect length
    for (loc in c(0.5*len, 0.75*len, 0.9*len)){ # band location (mean for dnorm)
      for (sca in c(1, 2, 5, 10, 20)){ # band scale (sd for dnorm)
        for (myn in c(3, 6, 11, 16)){ # number of levels
          i<-i+1
          results2[i,1]<-tot
          results2[i,2]<-len
          results2[i,3]<-loc
          results2[i,4]<-sca
          ## configure sampling
          int<-len/(myn-1) # interval
          results2[i,5]<-int
          results2[i,6]<-myn
          myx<-seq(0, len, int) # level locations
          ## sample
          myy<-tot*dnorm(myx, loc, sca) # level densities (ind/m2)
          dens<-int*(0.5*myy[1]+sum(myy[2:(myn-1)])+0.5*myy[myn]) # integrated density
          band<-qnorm(0.975, loc, sca)-qnorm(0.025, loc, sca) # 95% band width
          trunc<-(loc+0.5*band)-len # positive truncation if upper limit of 95% band > length
          results2[i,7]<-round(dens, 1) # integrated density
          results2[i,8]<-results2[i,7]-tot # bias
          results2[i,9]<-round((dens-tot)/tot, 3) # relative bias
          results2[i,10]<-round(mean(myy), 3) # mean level density
          results2[i,11]<-round(band, 1)
          results2[i,12]<-round(trunc, 1)
          results2[i,13]<-round(trunc/band, 3) # relative truncation
          results2[i,14]<-round(int/band, 3) # sampling frequency
        } # nlev
      } # sca
    } # loc
  } # len
} # tot
results2<-as.data.frame(results2)
names(results2)<-c("tot", "len", "loc", "sca", "int", "levels", "ldens", "bias", "relb", "adens", "band", "trunc", "relt", "freq")

# result 1: sampling frequency problem
# ------------------------------------
# similar to fixed interval simulations above

# remove truncation
freqs<-subset(results2, trunc<0)
# examine the effect of level interval and macrofauna band width on linear density
par(mfrow=c(2,2))
with(freqs, plot(ldens~levels, ylim=c(0,1500)))
abline(h=100, lty="dotted")
abline(h=200, lty="dotted")
abline(h=400, lty="dotted")
with(freqs, plot(relb~levels, ylim=c(-1,4)))
abline(h=0, lty="dotted")
with(freqs, plot(ldens~freq, xlab="int/band", xlim=c(0,3), ylim=c(0,1500)))
abline(h=100, lty="dotted")
abline(h=200, lty="dotted")
abline(h=400, lty="dotted")
abline(v=0.5, lty="dashed")
with(freqs, plot(relb~freq, xlab="int/band", xlim=c(0,3), ylim=c(-1,4)))
abline(h=0, lty="dotted")
abline(v=0.5, lty="dashed")
par(mfrow=c(1,1))

# result 2: truncation problem
# ----------------------------
# similar to fixed interval simulations above

# remove undersampling
truncs<-subset(results2, freq<=0.5)
# examine the effect of truncated distributions on linear density
par(mfrow=c(2,2))
with(truncs, plot(ldens~trunc))
abline(h=100, lty="dotted")
abline(h=200, lty="dotted")
abline(h=400, lty="dotted")
abline(v=0, lty="dashed")
with(truncs, plot(relb~trunc))
abline(h=0, lty="dotted")
with(truncs, plot(ldens~relt, xlab="trunc/band", xlim=c(-5,2)))
abline(h=100, lty="dotted")
abline(h=200, lty="dotted")
abline(h=400, lty="dotted")
abline(v=0, lty="dashed")
with(truncs, plot(relb~relt, xlab="trunc/band", xlim=c(-1,2)))
abline(h=0, lty="dotted")
abline(0,-1, lty="dashed")
text(0.7, -0.2, "y = -x")
par(mfrow=c(1,1))

# result 3: width problem
# -----------------------
# similar to fixed interval simulations above

# remove truncation and undersampling
best2<-subset(results2, trunc<0) # no truncation
best2<-subset(best2, freq<=0.5) # no undersampling
# fit a model to adens ~ len
amodel2<-lm(adens~tot*as.factor(len), best2)
summary(amodel2)
# plot results
best3<-subset(best2, levels==6)
with(best3, plot(adens~tot, ylim=c(0,16)))
best3<-subset(best2, levels==11)
with(best3, points(adens~tot, col="red"))
best3<-subset(best2, levels==16)
with(best3, points(adens~tot, col="blue"))
abline(coef(amodel2)[1], coef(amodel2)[2], lty="dashed") # 30 m
abline(coef(amodel2)[1]+coef(amodel2)[3], coef(amodel2)[2]+coef(amodel2)[5], lty="dashed") # 60 m
abline(coef(amodel2)[1]+coef(amodel2)[4], coef(amodel2)[2]+coef(amodel2)[6], lty="dashed") # 120 m

# --------------------------------------------
# simulation 3 (with negative binomial counts)
# --------------------------------------------
# the negative binomial model is useful for strongly aggregated counts
# the negative binomial variance = mean + (mean^2)/theta
# the parameter theta decribes the degree of clumping (overdispersion relative the poisson counts) and VARIES WITH QUADRAT SIZE, SPACING BETWEEN QUADRATS AND DIFFERENT SPECIES; pilot studies are needed to estimate theta (and the width of bands) for species of interest  
# here, theta = 1.6 is based on extensive sampling of the surf clam D. deltoides with 0.1 m2 quadrats with 5 paces between quadrats (56 samples of 7 quadrats from 15 beaches; unpub. data)
# simulation 2 is similar to simulation 1 plus replicate quadrats and multiple runs (replicate transects)

library(MASS) # for rnegbin

stacked<-array(dim=c(100, 3*3*3*5*6*3, 15))
for (run in 1:100){
  results<-array(dim=c(3*3*3*5*6*3, 15))
  i<-0 # index
  ## configure across transect abundance and distribution
  for (tot in c(100, 200, 400)){ # transect total (ind/m)
    for (len in c(30, 60, 120)) { # transect length
      for (loc in c(0.5*len, 0.75*len, 0.9*len)){ # band location (mean for dnorm)
        for (sca in c(1, 2, 5, 10, 20)){ # band scale (sd for dnorm)
          for (int in c(1, 3, 6, 10, 20, 40)){ # transect level interval
            for (rep in c(1, 3, 10)){ # number of replicate quadrats per level
              i<-i+1
              results[i,1]<-tot
              results[i,2]<-len
              results[i,3]<-loc
              results[i,4]<-sca
              results[i,5]<-int
              results[i,6]<-rep
              ## configure sampling
              myx<-seq(0, len, int) # level locations
              # check that len is divisible by int
              # forbidden combinations are (len, int) = (30, 20), (30, 40), (60, 40)
              if (max(myx)==len){
                ## sample
                myn<-length(myx) # number of levels
                # dnorm simulates the across shore distribution
                smooth<-tot*dnorm(myx, loc, sca) # smooth density
                myy<-array(dim=myn) # empty array
                for (j in 1:myn){
                  # rnegbin simulates quadrat counts with mean from smooth density
                  # level density is the sum of quadrat counts divided by quadrat area 
                  myy[j]<-sum(rnegbin(rep, 0.1*smooth[j], 1.6))/(rep*0.1)
                }
                dens<-int*(0.5*myy[1]+sum(myy[2:(myn-1)])+0.5*myy[myn]) # integrated density
                band<-qnorm(0.975, loc, sca)-qnorm(0.025, loc, sca) # 95% smooth band width
                trunc<-(loc+0.5*band)-len # positive truncation if upper limit of 95% band > length
                results[i,7]<-myn
                results[i,8]<-round(dens, 1) # integrated density
                results[i,9]<-results[i,8]-tot # bias
                results[i,10]<-round((dens-tot)/tot, 3) # relative bias
                results[i,11]<-round(mean(myy), 3) # mean level density
                results[i,12]<-round(band, 1)
                results[i,13]<-round(trunc, 1)
                results[i,14]<-round(trunc/band, 3) # relative truncation
                results[i,15]<-round(int/band, 3) # sampling frequency
              } # endif
            } # rep
          } # int
        } # sca
      } # loc
    } # len
  } # tot
  stacked[run, , ]<-results
} # run

# result 4: noise problem
# -----------------------
# noisy level counts result in noisy densities
# noise is reduced by sampling more quadrats, by sampling more replicate quadrats per level or more replicate transects
# the critical sampling frequency (see result 1) is still apparent when there is "stacking" of quadrats within levels

# stacked data
## meanstack n=100
# stacking equals 100 * rep quadrats = 100, 300, 1000 (enormous sampling effort)
meanstack100<-apply(stacked, c(2, 3), mean)
meanstack100<-as.data.frame(meanstack100)
names(meanstack100)<-c("tot", "len", "loc", "sca", "int", "rep", "levels", "ldens", "bias", "relb", "adens", "band", "trunc", "relt", "freq")
freqs100<-subset(meanstack100, trunc<=0) # no truncation
## meanstack n=10
# stacking equals 10 * rep quadrats = 10, 30, 100 (moderate-large sampling effort)
subsample<-sample(1:100, 10)
substack<-stacked[subsample, , ]
meanstack10<-apply(substack, c(2, 3), mean)
meanstack10<-as.data.frame(meanstack10)
names(meanstack10)<-c("tot", "len", "loc", "sca", "int", "rep", "levels", "ldens", "bias", "relb", "adens", "band", "trunc", "relt", "freq")
freqs10<-subset(meanstack10, trunc<=0) # no truncation
## meanstack n=1
# stacking equals 1 * rep quadrats = 1, 3, 10 (small-moderate sampling effort)
subsample<-sample(1:100, 1)
meanstack1a<-stacked[subsample, , ]
meanstack1a<-as.data.frame(meanstack1a)
names(meanstack1a)<-c("tot", "len", "loc", "sca", "int", "rep", "levels", "ldens", "bias", "relb", "adens", "band", "trunc", "relt", "freq")
freqs1a<-subset(meanstack1a, trunc<=0) # no truncation
## meanstack n=1 (noise is random and runs vary)
subsample<-sample(1:100, 1)
meanstack1b<-stacked[subsample, , ]
meanstack1b<-as.data.frame(meanstack1b)
names(meanstack1b)<-c("tot", "len", "loc", "sca", "int", "rep", "levels", "ldens", "bias", "relb", "adens", "band", "trunc", "relt", "freq")
freqs1b<-subset(meanstack1b, trunc<=0) # no truncation

# examine noise levels for different levels of stacking
par(mfrow=c(2,2))
# 100 transects
with(freqs100, plot(ldens~freq, xlab="int/band", xlim=c(0,3), ylim=c(0,1000)))
abline(h=100, lty="dotted")
abline(h=200, lty="dotted")
abline(h=400, lty="dotted")
abline(v=0.5, lty="dashed")
# 10 transects
with(freqs10, plot(ldens~freq, xlab="int/band", xlim=c(0,3), ylim=c(0,1000)))
abline(h=100, lty="dotted")
abline(h=200, lty="dotted")
abline(h=400, lty="dotted")
abline(v=0.5, lty="dashed")
# 1 transect
with(freqs1a, plot(ldens~freq, xlab="int/band", xlim=c(0,3), ylim=c(0,1000)))
abline(h=100, lty="dotted")
abline(h=200, lty="dotted")
abline(h=400, lty="dotted")
abline(v=0.5, lty="dashed")
# 1 transect
with(freqs1b, plot(ldens~freq, xlab="int/band", xlim=c(0,3), ylim=c(0,1000)))
abline(h=100, lty="dotted")
abline(h=200, lty="dotted")
abline(h=400, lty="dotted")
abline(v=0.5, lty="dashed")
par(mfrow=c(1,1))

# examine noise levels for n > 1 replicates
par(mfrow=c(2,2))
# 100 transects
freqs100r=subset(freqs100, rep>1)
with(freqs100r, plot(ldens~freq, xlab="int/band", xlim=c(0,3), ylim=c(0,1000)))
abline(h=100, lty="dotted")
abline(h=200, lty="dotted")
abline(h=400, lty="dotted")
abline(v=0.5, lty="dashed")
# 10 transects
freqs10r=subset(freqs10, rep>1)
with(freqs10r, plot(ldens~freq, xlab="int/band", xlim=c(0,3), ylim=c(0,1000)))
abline(h=100, lty="dotted")
abline(h=200, lty="dotted")
abline(h=400, lty="dotted")
abline(v=0.5, lty="dashed")
# 1 transect
freqs1ar=subset(freqs1a, rep>1)
with(freqs1ar, plot(ldens~freq, xlab="int/band", xlim=c(0,3), ylim=c(0,1000)))
abline(h=100, lty="dotted")
abline(h=200, lty="dotted")
abline(h=400, lty="dotted")
abline(v=0.5, lty="dashed")
# 1 transect
freqs1br=subset(freqs1b, rep>1)
with(freqs1br, plot(ldens~freq, xlab="int/band", xlim=c(0,3), ylim=c(0,1000)))
abline(h=100, lty="dotted")
abline(h=200, lty="dotted")
abline(h=400, lty="dotted")
abline(v=0.5, lty="dashed")
par(mfrow=c(1,1))

# result 5: smaller intervals reduce noise
# ----------------------------------------
# the precision of mean linear densities is similar whether replication of quadrats is along-shore (within levels) or across-shore (and closely spaced at 5 paces) 

## filtered data
# remove truncation
myselect<-stacked[,,13]<=0 # each of the 100 sims are identical for trunc
freqs<-stacked[myselect[,1], myselect[1,], ]
# remove undersampling 
myselect2<-freqs[,,15]<=0.5
freqs2<-freqs[myselect2[,1], myselect2[1,], ]
# remove NAs
myselect3<-(is.na(freqs2[,,14])==FALSE)
freqs3<-freqs2[myselect3[,1], myselect3[1,], ]
dim(freqs)
dim(freqs2)
dim(freqs3)
## stacked data, mean and variance
meanstack<-apply(freqs3, c(2, 3), mean) # mean of 100 sims
varstack<-apply(freqs3, c(2, 3), var) # var of 100 sims
meanstack<-as.data.frame(meanstack)
varstack<-as.data.frame(varstack)
names(meanstack)<-c("tot", "len", "loc", "sca", "int", "rep", "levels", "ldens", "bias", "relb", "adens", "band", "trunc", "relt", "freq")
names(varstack)<-c("tot", "len", "loc", "sca", "int", "rep", "levels", "ldens", "bias", "relb", "adens", "band", "trunc", "relt", "freq")
head(meanstack)
head(varstack)
# bind relevant variables
mydata<-cbind(meanstack, varstack[,8:10])
names(mydata)[16:18]<-c("vdens", "vbias", "vrelb")
head(mydata)
unique(mydata$int)
unique(mydata$rep)
## matched data for 1 replicate quadrat at 1 m intervals and 3 replicates at 3 m
int1<-subset(mydata, int==1)
int1<-subset(int1, rep==1)
int1a<-subset(int1, sca>1) # because sca = 1 results in undersampling for int = 3
rep3<-subset(mydata, int==3)
rep3<-subset(rep3, rep==3)
# check simulations match
dim(int1a)
dim(rep3)
int1a[,1:4]-rep3[,1:4]
# efforts are
int1a$rep*int1a$levels
rep3$rep*rep3$levels
# the difference in effort is consistently -2 quadrats (slightly greater effort with 3 replicates at 3 m)
int1a$rep*int1a$levels - rep3$rep*rep3$levels
## matched data for 1 replicate quadrat at 1 m intervals and 10 replicates at 10 m
int1<-subset(mydata, int==1)
int1<-subset(int1, rep==1)
int1b<-subset(int1, sca>5) # because sca = 5 results in undersampling for int = 10
rep10<-subset(mydata, int==10)
rep10<-subset(rep10, rep==10)
# check simulations match
dim(int1b)
dim(rep10)
int1b[,1:4]-rep10[,1:4]
# efforts are
int1b$rep*int1b$levels
rep10$rep*rep10$levels
# the difference in effort is consistently -9 quadrats (greater effort with 10 replicates at 10 m)
int1b$rep*int1b$levels - rep10$rep*rep10$levels

## compare 1 at 1 m to 3 at 3 m and 1 at 1 m to 10 at 10 m
par(mfrow=c(2,3))
plot(int1a$ldens, rep3$ldens) # densities
abline(0, 1, lty="dotted")
plot(int1a$vdens, rep3$vdens) # variances
abline(0, 1, lty="dotted")
plot(sqrt(int1a$vdens)/int1a$ldens, sqrt(rep3$vdens)/rep3$ldens) # coefficients of variation
abline(0, 1, lty="dotted")
plot(int1b$ldens, rep10$ldens) # densities
abline(0, 1, lty="dotted")
plot(int1b$vdens, rep10$vdens) # variances
abline(0, 1, lty="dotted")
plot(sqrt(int1b$vdens)/int1b$ldens, sqrt(rep10$vdens)/rep10$ldens) # coefficients of variation
abline(0, 1, lty="dotted")
par(mfrow=c(1,1))
